# Sonic Hedgehog is a member of the Hh/DD-peptidase family that spans the eukaryotic and bacterial domains of life

**DOI:** 10.1101/276295

**Authors:** Henk Roelink

## Abstract

Sonic Hedgehog (Shh) coordinates Zn^2+^ in a manner that resembles that of peptidases. The ability of Shh to undergo autoproteolytic processing is impaired in mutants that affect the Zn^2+^ coordination, while mutating residues essential for catalytic activity results in more stable forms of Shh. The residues involved in Zn^2+^ coordination in Shh are found mutated in some individuals with the congenital birth defect holoprosencephaly, demonstrating their importance in development. Highly conserved Shh domains are found a parts of some bacterial proteins that are members of the larger family of DD-peptidases, supporting the notion that Shh acts as a peptidase. Whereas this Hh/DD-peptidase motif is present in Hedgehog (Hh) proteins of nearly all animals, it is not present in *Drosophila* Hh, indicating that Hh signaling in fruit flies is derived, and perhaps not a good model for vertebrate Shh signaling. Sequence analysis of Hh proteins and their possible evolutionary precursors suggest that the evolution of modern Hh might have involved horizontal transfer of a bacterial gene coding of a Hh/DD-peptidase into a *Cnidarian* ancestor, recombining to give rise to modern Hh.

## The Sonic Hedgehog pro-protein gives rise to the mature ligand after an autoproteolytic cleavage event

The *Hedgehog* (*Hh*) gene was first identified in the now famous developmental *Drosophila melanogaster* screen performed by Christiane Nüsslein-Volhard and Eric Wieshaus in the late 1970s. The screen used a technique known as “saturation mutagenesis” to isolate genes involved in the formation of the Drosophila body plan [1]. *Hh* mutant larvae have a solid lawn of denticles on the cuticle rather than stripes of denticles on the anterior half of each segment, hence the name “hedgehog”. Like other segment polarity genes found in this screen, also *Hh* genes are widely conserved among animals, and mammals have three Hh paralogs (Sonic, Indian, and Desert Hedgehog) that, like in *Drosophila*, play central roles in development [2].

Sonic Hedgehog (Shh) coordinates a Zn^2+^ metal ion with residues H141, D148, and H183 (mouse numbering, Figure 1) typical for Zn^2+^ peptidases, [3] such as the bacterial peptidase thermolysin [4]. Like many peptidases, the Shh undergoes an intramolecular auto-processing reaction resulting in cleavage between G198 and C199. As a consequence of this cleavage event the N-terminal product of this cleavage (ShhN), is modified with cholesterol [5,6]. Subsequently Shh is modified by an N-terminal acylation [7–9], rendering ShhN_chol_ obligatory membrane bound. Secretion of this form of Shh requires Disp1and Scube2 [10] and ADAM-type metalloproteases, yielding a form that is stripped of its lipid modifications active in signaling [11,12] (Figure 1). Whereas ShhN harbors the Zn^2+^ peptidase motif, the carboxyterminal domain has similarities to self-splicing bacterial inteins [13]. These inteins typically cleave before a Cysteine residue through the resolution of a thio-ester intermediate [6]. The G198/C199 site of cleavage is consistent with this idea. Single amino acid changes in C-terminal domains can prevent auto-processing, resulting in the perdurance of the Shh pro-protein [14]. Similarly, mutations of the residues that directly mediate Zn^2+^ coordination prevent the autoproteolytic processing of the Shh [15] and thermolysin pro-proteins [16]. There thus appear to be structural requirements in both the ShhC and ShhN domains for auto-processing to proceed, perhaps indicating that overlapping/complementary endopeptidase activities associated with the ShhN and −C domains are involved in the autocatalytic processing. The inherent endopeptidase activity of Shh is coupled to the addition of a cholesterol moiety to ShhN (ShhNChol)), which affects distribution of the ligand [17]. Interestingly, many of the Zn^2+^ peptidase catalytic residues are not required for signaling by ShhN [18], and consequently the Zn^2+^ coordination domain of Shh has been referred to as the “pseudo active” site [19,20]. Nevertheless, the importance of the Zn^2+^ coordination domain has become apparent as mutations of this domain are associated with a congenital malformation, demonstrating a role in development.

**Figure 1.**
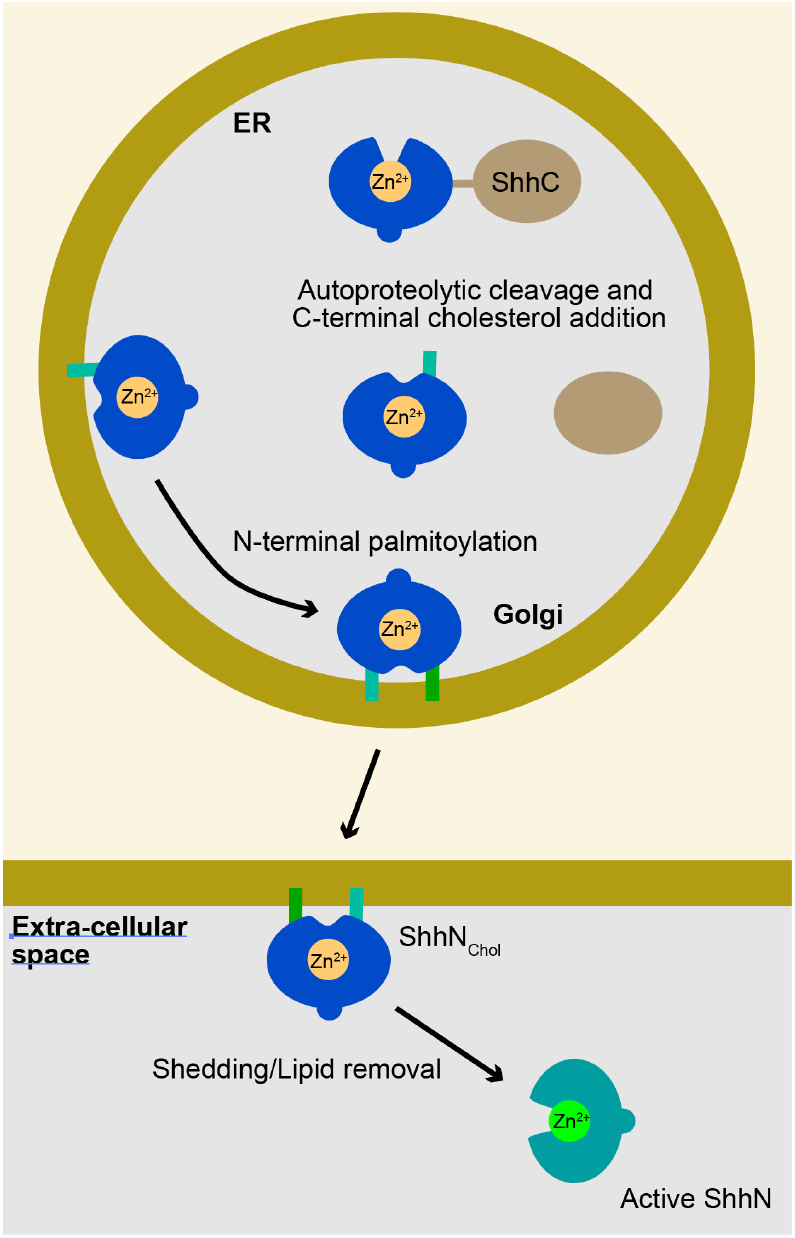
Diagram of the Shh processing/maturation steps. Shh is translated into a pro-protein, consisting of a ShhN and ShhC domain, that undergoes an autoproteolytic processing event prior to entry into the Golgi. This results in the attachment of a cholesterol moiety the ShhN domain, likely shielding the Zn^2+^ coordination domain. In the Golgi, ShhN_chol_ is further lipidated by a palmitoyl chain (green bars), further forcing its membrane association. Release and shedding is a regulated process involving Disp1, Scube2 and ADAM metalloproteases, resulting in the release of Shh devoid of its lipid moieties, and its Zn^2+^ coordination domain exposed.

## Both the N- and C-terminal domains of Shh are the targets for point mutations found in holoprosencephaly

SHH mutations are commonly found in holoprosencephaly a congenital syndrome that can be caused by aberrant SHH signaling [21–23] (Figure 2C). Single amino acid substitutions can be found in both the N- and C-domains of the SHH pro-protein, but are more prevalent in SHHN (40/181 in N, 38/266 in C, Z=−3.6, *p*=3.2×10^−4^). Two of the Zn^2+^ coordination residues (H140 and D147 (Figure 2C, blue), have been found mutated in holoprosencephalic individuals, indicating that they are required for normal SHH function, consistent with the notion that the putative peptidase activity of Shh is important for signaling. The D148 equivalent is not conserved in *Drosophila* Hh, indicating it is not required for binding to Ptch. Traiffort *et al*. showed that SHH-H140P failed to undergo auto-processing, and was detected only as the SHH pro-protein [15]. This indicates that the perdurance of the SHH pro-protein might contribute to holoprosencephaly. It further shows that the correct Zn^2+^ coordination is necessary for processing the Shh pro-protein into ShhN_Chol_. The face of ShhN opposite of the Zn^2+^ coordination domain is dominated by a large α–Helix (Figure 2A). This helix is enriched in point mutations found in holoprosencephalic individuals (Figure 2C, dark green). Two tested mutations, SHH-W117G and W117R, were unable to undergo auto-processing [15], further emphasizing structural requirements of the N-domain in auto-processing. Similarly, several mutations in the C-terminal domain prevent processing [14,15], emphasizing the central role that this domain play in processing the Shh pro-protein. The SHH mutations found in holoprosencephaly that thus likely affect SHH function indicate critical roles for both the N-terminal and C-terminal domains in auto-processing leaving the precise mechanisms and events by which the SHH pro-protein matures unresolved.

**Figure 2.**
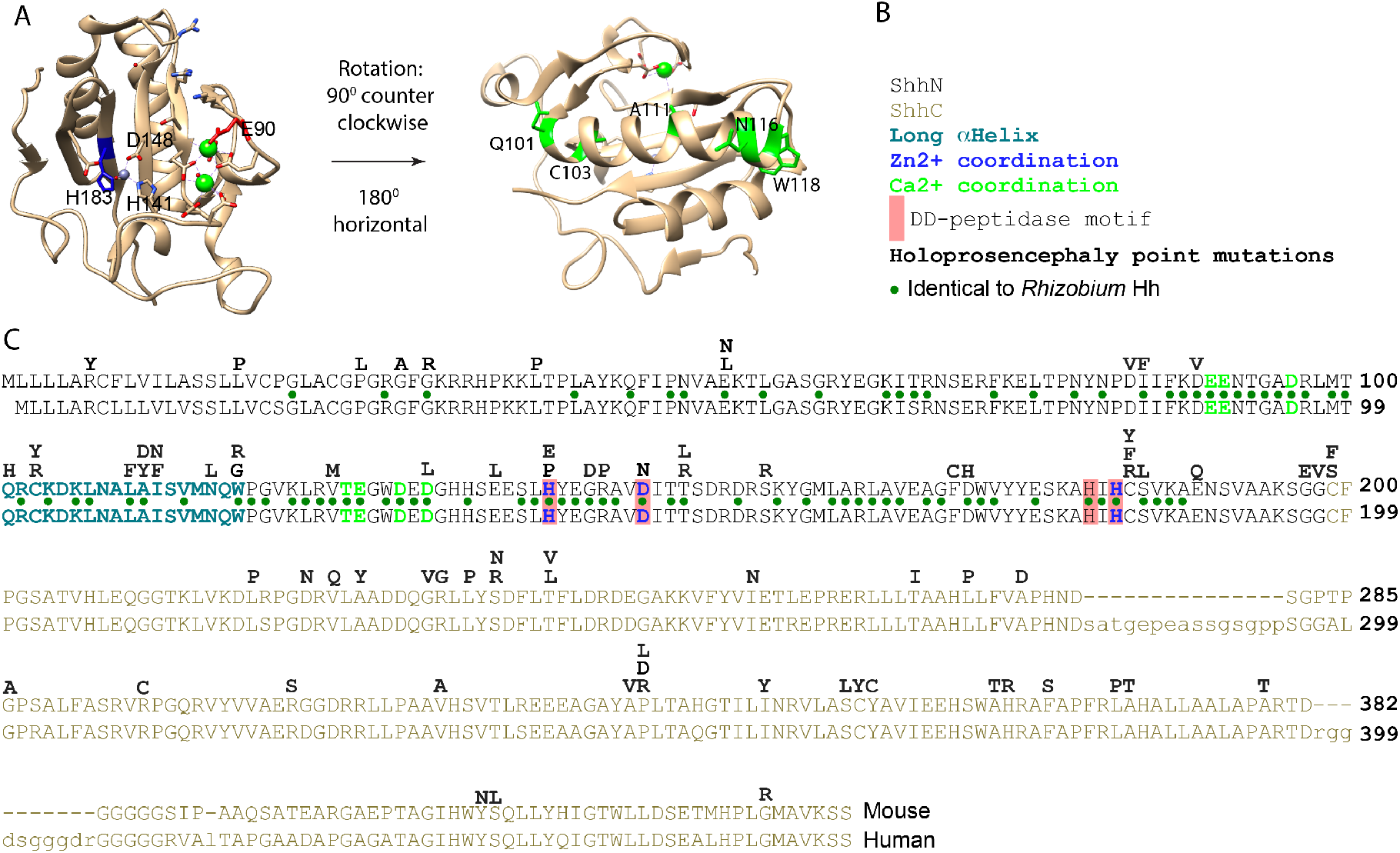
Salient features of the Shh protein. (A) Crystal structure of ShhN. The Zn^2+^ (steel) and Ca^2+^ (green) coordination domains indicated. Residues in the large α-helix mutated in Holoprosencephaly are labeled in green. (B) Legend. (C) Lineup of mouse Shh (top) and human (bottom) SHH. Point mutations resulting in single amino acid changes in SHH found in holoprosencephalic individuals are above the lineup. Residues mediating Ca^2+^ and Zn^2+^ coordination are labeled in blue and green respectively. Green dots indicate conserved residues between *Rhizobium* Hh and mouse/human SHH.

## Shh has all hallmarks of a DD-peptidase

The notion that Hhs are pseudo-proteases is primarily based on studies claiming that the Zn^2+^ coordination site is required to maintain the correct and stable Shh structure, and for Shh binding to Ptch1 [24], but does not mediate protease activity [25]. This rejection of Shh protease function were based one experiments with negative outcomes, based on simple biochemical assays using artificial peptide substrates, common peptidase inhibitors, and *E.coli* expressed non-lipidated ShhN mutants that were not derived from the Shh pro-protein. Under such experimental conditions, even testing established proteases would possibly fail to reveal their proteolytic activitiesy. A possibly more useful approach would be to more precisely determine the effects of Robotnikinin, a small molecule that binds to the Shh Zn^2+^ coordinating domain [26] at high nanomolar concentrations [27], in cells that lack Ptch function.

In Zn^2+^ peptidases, the E177 equivalent abstracts a proton from the catalytic water at the Zn^2+^ coordination domain, which is followed by a nucleophilic attack of the OH-on the peptide backbone. Shh-E177A is, therefore, predicted to be impaired for the intrinsic Zn^2+^ peptidase activity. Analysis of this mutant in has revealed two interesting properties. First, Shh-E177A is unable to mediate signaling from the notochord to the overlying neural tube (*in trans*), but is more capable than Shh to induce the Hh response when expressed in the developing neural tube (likely *in cis*) [28]. Second, purified ShhN-E177A is more stable in solution than ShhN, indicating a cannibalistic peptidase activity that is intrinsic to ShhN. This activity is inhibited by the binding of one or two Ca^2+^ ions [4] to ShhN (Figure 2A), using a binding motif that is conserved among Hh proteins, and present in thermolysin. It appears that interfering with the putative Zn^2+^ peptidase activity either via the Zn^2+^ coordination domain or E177 has negative consequences for Shh signaling during development, indicating a role for this peptidase activity-associated residue for normal Shh function.

The conservation of the Zn^2+^ coordinating, Ca^2+^ coordinating and other catalytic residues throughout evolution supports the possibility that Hhs are not pseudo-proteases but, using the properly matured form with the endogenous substrate, may indeed act as peptidases. Furthermore, structural analysis of interactions between Shh and the Hh interacting protein (Hhip) show that they resemble molecular interactions between matrix metalloproteases (MMPs) and their tissue inhibitor of metalloproteinase (TIMP). In both cases a loop present in Hhip and TIMP interacts closely with the ShhN-coordinated Zn^2+^ thus blocking catalytic activity. This striking similarity between unrelated protease/inhibitor pairs further supports the notion that ShhN is a Zn^2+^ peptidase: not only is an active site present, but interactions of this site with Hhip and possibly Ptch1 also resemble established metalloprotease/inhibitor interactions [19].

Typical for a protease active site, the Zn^2+^ in ShhN sits at the bottom of a cleft and is exposed to solvent, and not hidden inside of the molecule. This configuration that is conserved in the mature form of well-characterized protease thermolysin [29]. ShhN homologs in bacteria are characterized by a conserved Zn^2+^ coordination motif, that defines a family of prokaryotic proteins that are characterized by the DD-peptidase fold [30]. The signature peptidase fold is a central, five-stranded antiparallel β-sheet, separating the Zn^2+^ coordination domain from several α-helices, as found in ShhN (Figure 2A). Members of this family include murein endopeptidase (penicillin resistant enzymes) and peptidase M15 (bacterial D-alanyl-D-alanine carboxypeptidases (DD-peptidase), the target for penicillin) [30]. Both these peptidases play critical roles in bacterial cell wall modification. Shh and peptidase M15 share the two Histidine residues and an Aspartic acid residue that mediate Zn^2+^ coordination (H141 D148, H183 for Shh). The overall Zn^2+^ coordination motif is also found in lysostaphin, another peptidase that cleaves peptide bonds in bacterial peptidoglycan, and is referred to as the “LAS” (**L**ysostaphin, D-**a**lanyl-D-alanine carboxypeptidase, **S**hh) arrangement [31]. DD-peptidases characteristically contain a H-X(6)-D and H-X-H motif that coordinates Zn^2+^ in an accessible catalytic cleft. By all these criteria Shh is a member of this class of peptidases [30], further supporting the notion that Shh can function as a protease, perhaps even targeting a glycoprotein. Characterized bacterial DD-peptidases use a catalytic mechanism in which the acyl-linked peptide-enzyme intermediate is resolved by a nucleophilic attack of the adjacent peptidoglycan chain, thus cross-linking the peptidoglycan chains in the bacterial cell wall, and it remains to be determined if Shh has a similar activity.

## Several bacterial species have highly conserved HhN domains

Remarkably, several species of bacteria carry a highly conserved Hedgehog proteins of unknown function (Figure 3, 4, Appendix A). An about 145 amino acid domain comprising the bulk of ShhN (Figure 3, green box) and containing the typical DD-peptidase motif has over 50% <u>identity</u> to some bacterial proteins (Figure 4B), and I’ll refer to this class of protein domains as bacHhs. All the residues involved in Zn^2+^ and Ca^2+^ coordination are identical, as is E177, a residue required for non-autonomous Shh signaling. In bacteria, the Hh/DD-peptidases domain is the C-terminal part of larger proteins. Homologs of the N-terminal domains of the bacterial Hh proteins can be found associated with different subtypes of Zn^2+^ peptidases throughout bacteria (Figure 4A). In a few bacterial species, including some *Rhizobium* and *Bradyrhizobium* species, the N-terminal part preceding the Hh/DD-peptidase domain is predicted to contain three transmembrane regions, placing the Hh domain in the periplasmic space (Figure 4A). In this configuration, the third transmembrane domain od *Rhizobium* bacHh lines up perfectly with the signal sequence of Shh. No clear homologs of the Hh receptors Patched and Smoothened are present in any of these bacteria, or their *Rhizobium Legume* hosts, further supporting the idea that bacHhs serve as peptidases, rather than as ligands. The similarity of ShhN to its bacterial counterparts is as high as to other distant metazoan Hhs, and higher than to the *Cnidarian* hedgling N-domain, and *Drosophila* HhN (Figure 4B). Given the high degree of identity between Shh and its bacterial counterparts, it is likely that they share a specific function. The relatively small number of sequenced bacteria with a highly conserved HhN domain (I found about a dozen) do not share any obvious characteristics or ecological niches, and the precise function of this bacHh peptidase activity remains unknown.

**Figure 3.**
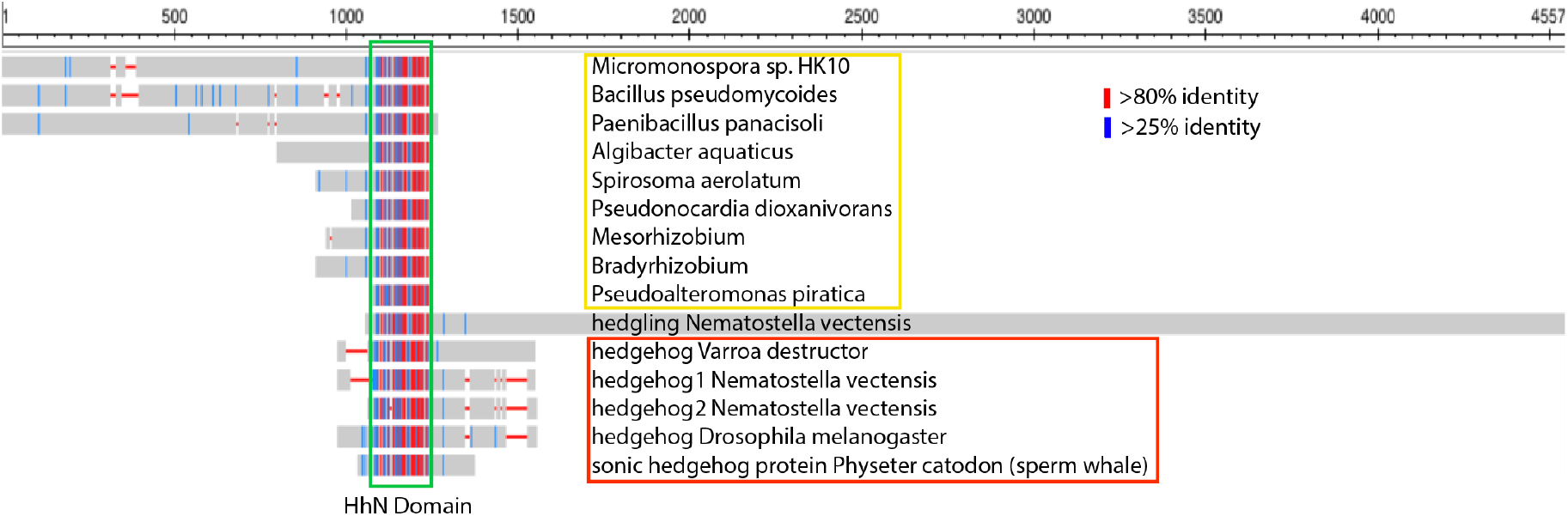
HhN domains are present in Bacteria. Lineup of several hypothetical proteins in bacteria, Hedgling and metazoan Hhs. The conserved domain (HhN, green box, about 175 residues) is flanked by other sequences in the various species. In bacteria (yellow box) the HhN domain is the C-terminal end of the hypothetical proteins. The HhN domain is the N-terminal part of hedgling and Hhs (red box). Blue and red lines indicate medium and high levels of conservation.

**Figure 4.**
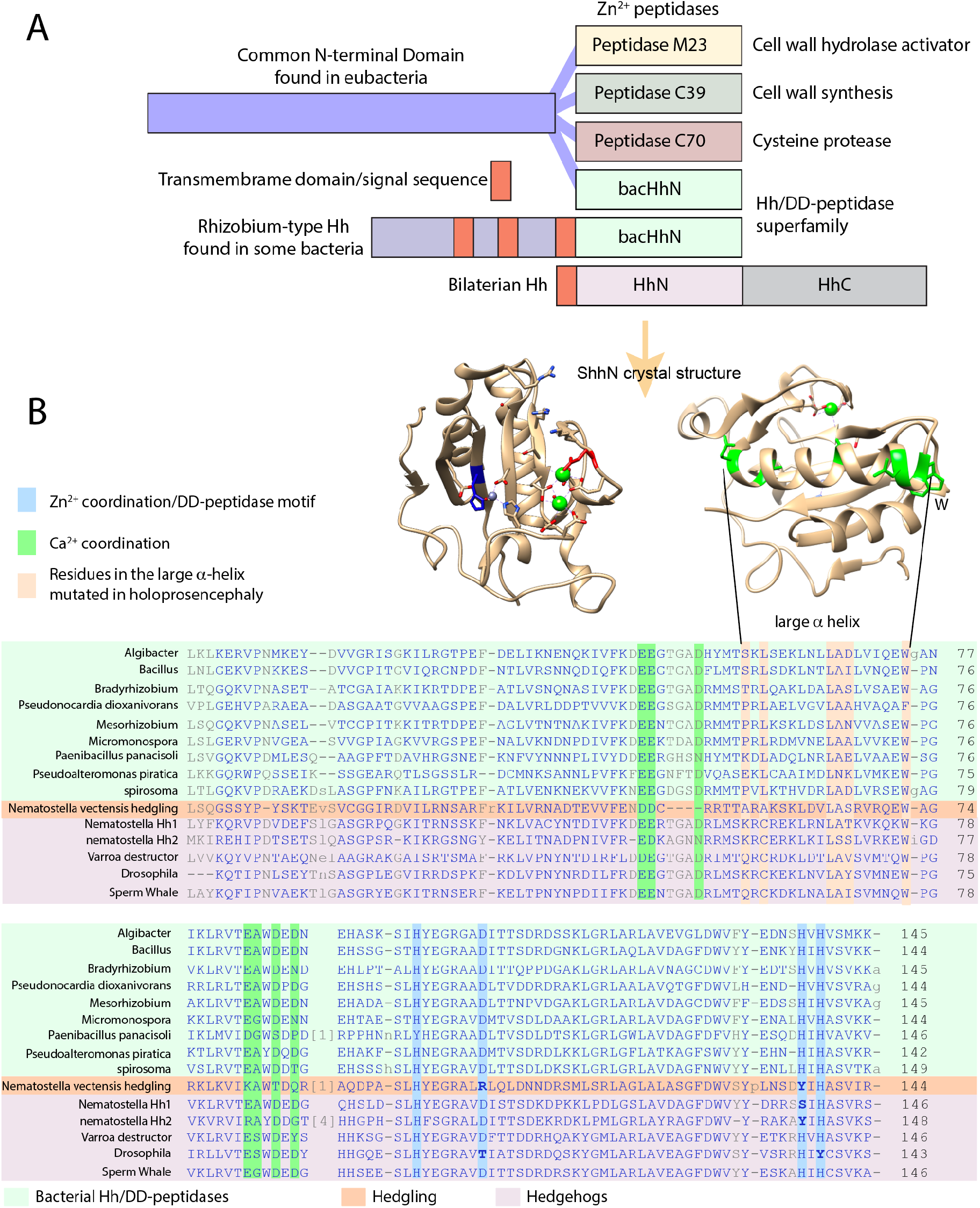
The Hh/DD-peptidase family. (A) Diagram of the structure of the DD-peptidases in bacteria and animals. In bacteria the peptidases are the C-terminal domain of larger proteins. In all cases the peptidase domain is predicted to be located in the periplasmic space. In *Rhizobium* Hh the third transmembrane domain (red) is at the same position as the signal sequence in metazoan Hh (red). (B) Lineup of the Hh/DD-peptidase domains of bacteria (green background), Hedgling (salmon background) and Hhs (purple background). Mutations in the Zn^2+^ coordinating/DD-peptidase motif defining residues in Hedgling and *Drosophila* Hh are indicated in bold. Blue columns indicate residues involved in Zn^2+^ coordination and define the DD-peptidase motif, green columns indicate residues involved in Ca2+ binding. *Varroa* is typical for all *non-Drosophilid* insects.

## What are the possible substrates for the Shh peptidase activity?

The general properties of bacterial DD-peptidases and lysostaphins as modifiers of the bacterial cell wall is likely shared with the bacHhs. The peptidoglycans that are a major component of the bacterial periplasmic space have some similarities to the proteoglycans that are common in the insect and vertebrate extracellular matrix. Both bacterial cell wall peptidoglycans and animal matrix proteoglycans are large molecules in which polypeptides are covalently attached to chains of glycans. In particular, the matrix heparin sulfate proteoglycans (HSPGs) bind Shh and can both negatively and positively affect the Shh response [32–34]. Furthermore, mutations in *ext* genes that code for glycosyltransferases that catalyze glycosaminoglycan addition to the core protein, disrupt Hh signaling in vertebrates [35] and insects [36]. It is thus possible that functional conservation between bacHhs and Shh is reflected in the ability of Shh to cleave or modify proteoglycans, thus affecting the Shh response or distribution, independent of binding to the canonical receptors.

Although any Shh antagonist could be a possible target for Shh peptidase activity, both Hhip and Ptch1 are unlikely substrates, as they have characteristics of metalloprotease inhibitors [19].

## *Drosophila* Hh is not a member of the Hh/DD-peptidase family

*Drosophila* Hh (dmHh) has been the guide molecule for all Hh signaling since its discovery as necessary for embryogenesis [1]. However, dmHh is unusual in that is does not have the core sequences H-X(6)-D, H-X-H that define the Hh/DD-peptidase motif, but instead dmHh has H-X(6)-**T**, H-X-**Y**. Furthermore, the catalytically important E177 (mouse numbering) is mutated in dmHh into a Valine residue. The absence of the Hh/DD-peptidase motif is unique to all sequenced *Drosophilids*; searching with the *Drosophila* sequence that encompasses the DD-peptidase motif (ESL**H**YEGRAV**T**IATSDRDQSKYGMLARLAVEAGFDWVSY**V**SRR**H**I**Y**CSVKS) places *Drosophilids* as an outgroup to all other arthropods and deuterostomes. *Drosophilids* are the outlier because Hh proteins in all other protostomes and deuterostomes retain the conserved Hh/DD-peptidase Zn^2+^ coordination motif. Searching with a hypothetical *dmHh* peptide that conforms with the Hh/DD-peptidase motif and E177 does no longer uniquely group with the *Drosophilids*, and now has similar homology to several vertebrate and insect Hhs. Given the high degree of HhN conservation between bacteria and eukaryotes (Figure 3), the most plausible explanation is that *Drosophilids* have lost some of the activities associated with the ancestral Zn^2+^ coordination domain. The loss of Zn^2+^ coordinating residues in conjunction with the E177 homolog is no surprise in the light of the likely loss of peptidase activity in *Drosophila* Hh. The observation that vertebrate Shh is active in *Drosophila* embryos [37], but *Drosophila Hh* is not active in vertebrates, further supports the notion that some aspects of Hh signaling present in most animals are lost in fruit flies, but that both proteins can bind to Ptch. These observations question to what extent the lessons learned in *Drosophila* embryos regarding Hh processing and signaling can be directly applied to vertebrates, or even other arthropods, as all these animals maintain an intact Hh/DD-peptidase domain. Study of Hh signaling in another insect, such a *Tribolium* [38] might help resolve how *Drosophila* Hh signaling is impacted by the loss of the DD-peptidase motif in the ligand.

## The presence of bacHhs suggest an alternative hypothesis regarding the evolution of modern Hh

*Hh* genes are present in *Cnidarians* (corals and jellyfish), but not in sponges and protozoa [39]. However, another class of proteins that contain an N-terminal HhN-like domain are the Hedgling proteins that can be found in *Cnidarians*, sponges and *Choanoflagellates* [40], which are protozoans. In these Hedgling proteins the HhN domain is followed by a large C-terminal domain that is related to cell adhesion molecules. A plausible evolutionary path is that the HhN domain of Hedgling was recombined to a Hog domain that is present in most genomes (Figure 5 green arrows). However, the HhN domain of Hedgling does not contain a complete Hh/DD-peptidase motif, whereas it is likely that its putative ancestral form in the last universal common ancestor (LUCA) had an intact Hh/DD-peptidase domain. Although this evolutionary path from a LUCA DD-peptidase via Hedgling to Hh could conceivably occur without horizontal gene transfer (Figure 5 green arrows), it does require several recombination events. The first event would entail the recombination of the bacterial-like Hh domain from a DD-peptidase to form the Hedgling protein, the second event would recombine the HhN domain of Hedgling into modern Hh. Following this potential evolutionary path the Hh/DD-peptidase motif, is lost in Hedgling, and reestablished in Hh.

**Figure 5.**
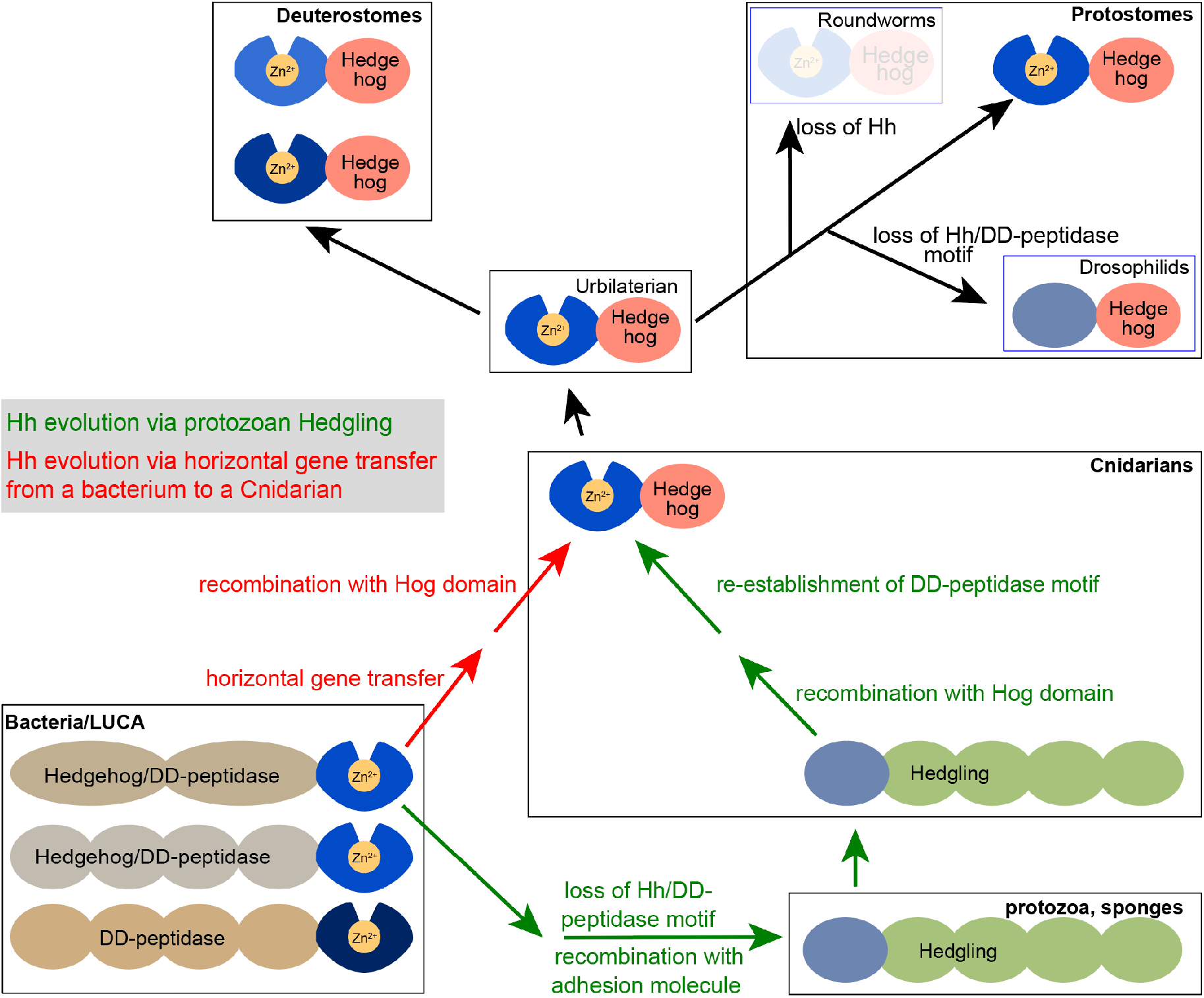
Models for the evolution of modern Hh proteins. The presence of both Hhs and Hedglings in Cnidarians support the model that the HhN of Hh domain arose via a recombination of the Hedgling N-domain. As Hedgling is present in at least some protozoa, it is plausible that is was derived from a DD-peptidase in the last universal common ancestor (LUCA). Alternatively, a Hh/DD-peptidase could have entered a Cnidarian ancestor from a bacterium via horizontal gene transfer, never losing the Zn^2+^ coordination in the process. Modern Hh was present in the Urbilaterian (the last common ancestor of protostomes and deuterostomes) and was retained in most of its offspring. The loss of the Hh/DD-peptidase motif in *Drosophilids* is derived, as is the loss of all Hh in some bilaterian protostomes (such as roundworms).

The presence of highly conserved HhN protein domains in bacteria suggests an alternate hypothesis, in which horizontal *bacHh/IDD-peptidase* gene transfer from a bacterium to a *Cnidarian* ancestor was followed by a recombination of the HhN domain with a *Cnidarian Hog* domain, resulting in a modern *Hh* gene (Figure 5, red arrows). This possible evolutionary path requires only one recombination event, and does not involve the subsequent loss and re-establishment of the Hh/DD-peptidase domain as is indicated for the evolutionary path via Hedgling, and is thus parsimonious path. The higher conservation between ShhN and bacterial Hh/DD-peptidases than between ShhN and the N-domain of Hedgling lends further support to the horizontal gene transfer model.

## Conclusions

Although the Zn^2+^ coordination domain of Shh is often referred to as its “pseudo catalytic” domain [19,41,42], the remarkable similarity of most Hhs to bacterial Hh/DD-peptidases further supports the notion that Shh functions as a peptidase during development. Some mutations found in holoprosencephaly patients break the Hh/DD-peptidase motif, and negatively affect Hh signaling possibly by preventing autoproteolytic cleavage, indicating that an intrinsic Zn^2+^ peptidase activity of Shh is critical to its function. The lack of the Hh/DD-peptidase domain in *Drosophila* Hh demonstrates that it is evolutionary derived, and perhaps not the best model for Hh signaling in animals with the ancestral Hh/DD-peptidase motif, like humans and mice. Finally, the presence of HhN protein domains in some bacteria might support an alternate pathway for the evolution of Hh, via horizontal gene transfer from bacteria into am Urbilaterian ancestor, resulting in modern Hh that retains an ancestral peptidase activity.

## Appendix A

Accession numbers for the proteins used for the lineup in Figure 3.

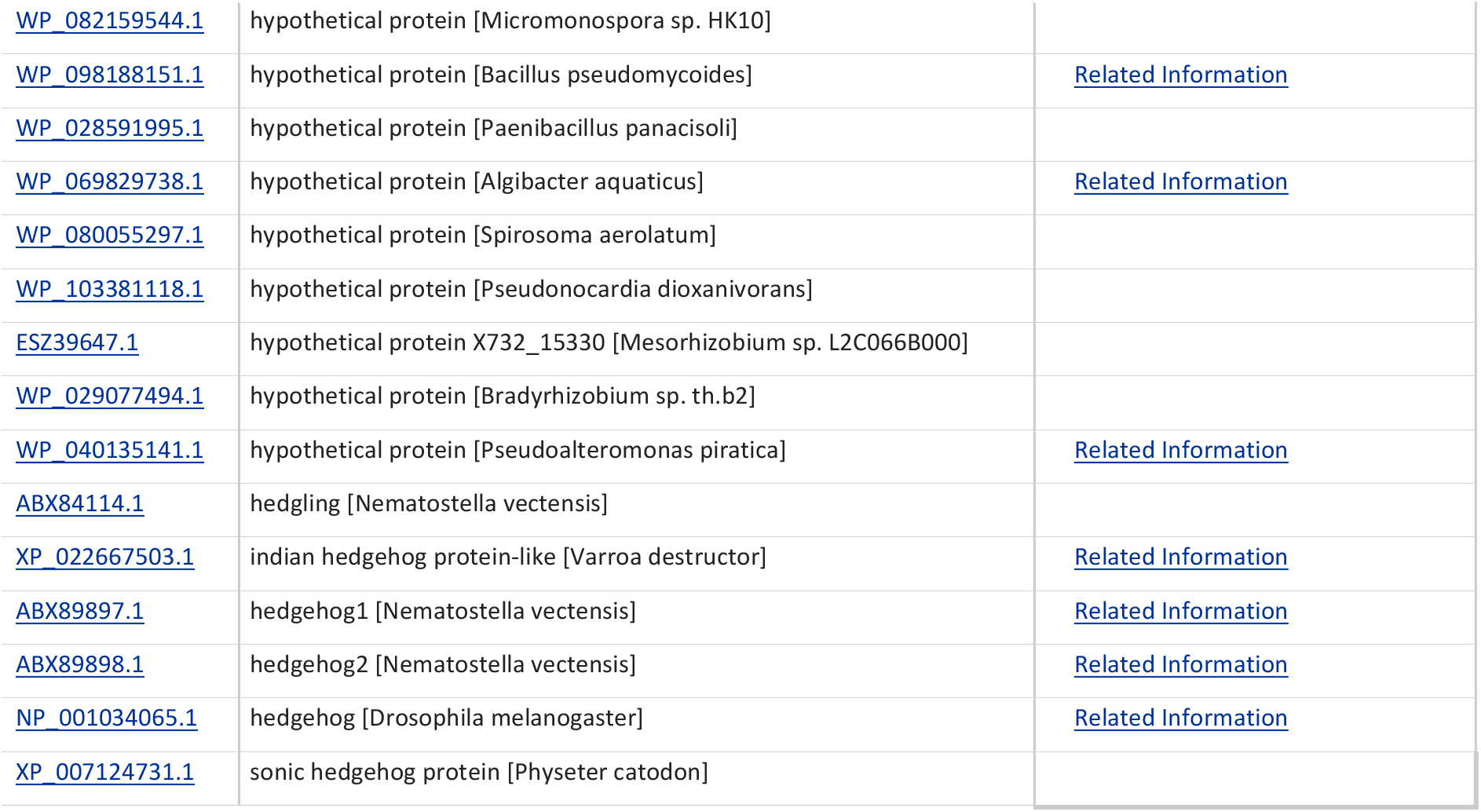

## Appendix B

BLAST Multiple alignment input (ncbi.nlm.nih.gov) for Hh/DD-peptidase domains used for the lineup in Figure 4

>Algibacter

LKLKERVPNMKEYDVVGRISGKILRGTPEFDELIKNENQKIVFKDEEGTGADHYMTSKLSEKLNLLADLVIQEWGANIKLRVTEAWDEDNEHASKSIHYEGRGADITTSDRDSSKLGRLARLAVEVGLDWVFYEDNSHVHVSMKK

>Bacillus

LNLGEKVPNKKESDVVGPITGVIQRGNPDFNTLVRSNNQDIQFKDEEGTGADFLMTSRLSDKLNTLAILVNQEWPNIKLRVTEAWDEDNEHSSGSTHYEGRAADITTSDRDGNKLGRLAQLAVDAGFDWVYYENKYHIHVSVKK

>Bradyrhizobium

LTQGQKVPNASETATCGAIAKKIKRTDPEFATLVSNQNASIVFKDEEGTGADRMMSTRLQAKLDALASLVSAEWAGVKLRVTEAWDENDEHLPTALHYEGRAADITTQPPDGAKLGRLARLAVNAGCDWVFYEDTSHVHVSVKKA

>Pseudonocardia dioxanivorans

VPLGEH

VPARAEADASGAATGVVAAGSPEFDALVRLDDPTVVVKDEEGSGADRMMTPRLAELVGVLAAHVAQAFPGRRLRLTEAWDPDGEHSHSSLHYEGRAADLTVDDRDRAKLGRLAALAVQTGFDWVLHENDHVHVSVRAG

>Mesorhizobium

LSQGQKVPNASELVTCGPITKKITRTDPEFAGLVTNTNAKIVFKDEENTGADRMMTPRLKSKLDSLANVVASEWPGAKLRVTEAWDEDNEHADASLHYEGRAADLTTNPVDGAKLGRLARLAVDAGCDWVFFEDSSHIHVSVKAG

>Micromonospora

LSLGERVPNVGEASVVGPIAGKVVRGSPEFNALVKNDNPDIVFKDEEKTDADRMMTPRLRDMVNELAALVVKEWPGKKLRVTEGWDENNEHTAESTHYEGRAVDMTVSDLDAAKLGRLARLAVDAGFDWVFYENALHVHASVKK

>Nematostella Hh1

LYFKQRVPDVDEFSLGASGRPQGKITRNSSKFNKLVACYNTDIVFKDEERTGADRLMSKRCREKLRNLATKVKQKWKGVKLRVTEAWDEDGQHSLDSLHYEGRAVDISTSDKDPKKLPDLGSLAVDAGFDWVYYDRRSSIHASVRS

>Nematostella Hh2

MKIREHIPDTSETSLQASGPSRKIKRGSNGYKELITNADPNIVFREDKAGNNRRMSKRCERKLKILSSLVRKEWIGDVKVRVIRAYDDGTSKKRHHGPHSLHFSGRALDITTSDEKRDKLPMLGRLAYRAGFDWVYRAKAYIHASVKS

>Nematostella vectensis hedgling

LSQGSSYPYSKTEVSVCGGIRDVILRNSARFRKILVRNADTEVVFENDDCRRTTARAKSKLDVLASRVRQEWAGRKLKVIKAWTDQRTAQDPASLHYEGRALRLQLDNNDRSMLSRLAGLALASGFDWVSYPLNSDYIHASVIR

>Paenibacillus panacisoli

LSVGQKVPDMLESQAAGPFTDAVHRGSNEFKNFVYNNNPLIVYDDEERGHSNHYMTKDLADQLNRLAELVAAE

WSGIKLMVIDGWSDPDVRPPHNNRLYHEGRAADLTVSDLDTSKLGRLGWLAVDAGFDFVHYESQDHIHVAVKV

>Pseudoalteromonas piratica

LKKGQRWPQSSEIKSSGEARQTLSGSSLRDCMNKSANNLPVFKFEEGNFTDVQASEKLCAAIMDLNKLVMKEWPGKTLRVTEAYDQDGEHAKFSLHNEGRAADMTVSDRDLKKLGRLGFLATKAGFSWVYYEHNHIHASVKR

>spirosoma

LTLGQKVPDRAEKDSLASGPFNKAILRGTPEFATLVENKNEKVVFKNEEGDGSDRMMTPVLKTHVDRLADLVRSEWGAGVSLRVTEAWDDTGEHSSSHSLHYEGRAVDLTTSDLDKSKLGRLGRLAVDAGFNWVYYENLLHIHASVTKA

>Varroa destructor Hh

LVVKQYVPNTAEQNEIAAGRAKGAISRTSMAFRKLVPNYNTDIRFLDDEGTGADRIMTQRCRDKLDTLAVSVMTQWPGVKLRVIESWDEYSHHKSGSLHYEGRAVDFTTDDRHQAKYGMLARLAVEAGFDWVYYETKRHVHASVKP

>Drosophila Hh

KQTIPNLSEYTNSASGPLEGVIRRDSPKFKDLVPNYNRDILFRDEEGTGADRLMSKRCKEKLNVLAYSVMNEWPGIRLLVTESWDEDYHHGQESLHYEGRAVTIATSDRDQSKYGMLARLAVEAGFDWVSYVSRRHIYCSVKS

>Sperm Whale Shh

LAYKQFIPNVAEKTLGASGRYEGKITRNSERFKELTPNYNPDIIFKDEENTGADRLMTQRCKDKLNALAISVMNQW

PGVKLRVTEGWDEDGHHSEESLHYEGRAVDITTSDRDRSKYGMLARLAVEAGFDWVYYESKAHIHCSVKA

